# Decoding semantics from dynamic brain activation patterns: From trials to task in EEG/MEG source space

**DOI:** 10.1101/2023.07.20.549326

**Authors:** Federica Magnabosco, Olaf Hauk

**Affiliations:** MRC Cognition and Brain Sciences Unit, University of Cambridge, UK

**Keywords:** semantic cognition, MVPA, decoding, task, semantics, visual word recognition

## Abstract

The temporal dynamics within the semantic brain network and its dependence on stimulus and task parameters are still not well understood. Here, we addressed this by decoding task as well as stimulus information from source-estimated EEG/MEG data. We presented the same visual word stimuli in a lexical decision (LD) and three semantic decision (SD) tasks. The meanings of the presented words varied across five semantic categories. Source space decoding was applied over time in five ROIs in the left hemisphere (Anterior and Posterior Temporal Lobe, Inferior Frontal Gyrus, Primary Visual Areas, and Angular Gyrus) and one in the right hemisphere (Anterior Temporal Lobe). Task decoding produced sustained significant effects in all ROIs from 50-100 ms, both when categorising tasks with different semantic demands (LD-SD) as well as for similar semantic tasks (SD-SD). In contrast, semantic word category could only be decoded in lATL, rATL, PTC and IFG, between 250-500 ms. Furthermore, we compared two approaches to source space decoding: Conventional ROI-by-ROI decoding and combined-ROI decoding with back-projected activation patterns. The former produced more reliable results for word-category decoding while the latter was more informative for task-decoding. This indicates that task effects are distributed across the whole semantic network while stimulus effects are more focal. Our results demonstrate that the semantic network is widely distributed but that bilateral anterior temporal lobes together with control regions are particularly relevant for the processing of semantic information.

**Significance Statement:** Most previous decoding analyses of EEG/MEG data have focussed on decoding performance over time in sensor space. Here for the first time we compared two approaches to source space decoding in order to reveal the spatio-temporal dynamics of both task and stimulus features in the semantic brain network. This revealed that even semantic tasks with similar task demands can be decoded across the network from early latencies, despite reliable differences in their evoked responses. Furthermore, stimulus features can be decoded in both tasks but only for a subset of ROIs and following the earliest task effects. These results inform current neuroscientific models of controlled semantic cognition.

## Introduction

Semantic cognition is the representation and processing of the acquired knowledge about the world. The semantic brain network that enables us to store, employ, manipulate, and generalise conceptual knowledge can be characterised along two dimensions: representation and control (Lambon Ralph et al., 2017). Bilateral anterior temporal lobes (ATLs) constitute a multimodal hub within the representation component, where modality-specific information is combined to represent coherent concepts (Patterson et al., 2007). However, context and task influence the type of information that is relevant in any particular moment, requiring a control system capable of manipulating and shaping the activations in the representation system (Jefferies, 2013). A recent meta-analysis indicated that the control system is mostly left-hemispheric and comprises inferior frontal gyrus (IFG) and posterior temporal cortex (PTC) (Jackson, 2021). However, little is known about the dynamics within and between the components of the semantic network. In this study, we are tapping into semantic brain dynamics by decoding task as well as stimulus features from source-estimated EEG/MEG data in a visual word recognition paradigm.

In neuroimaging research, EEG/MEG and fMRI datasets have traditionally been analysed with different decoding/multi-voxel-pattern-analysis approaches (Grootswagers et al., 2017). While fMRI studies often focus on *spatial information* of decoding accuracies across different cortical regions (e.g., Haxby et al., 2001; Huth et al., 2016), EEG/MEG studies have mostly focussed on the *time cours*e of decoding accuracy determined in sensor space (e.g., Cichy et al., 2014). However, ROI-wise decoding in source space is possible using appropriate distributed source estimation methods (e.g., Kietzmann et al., 2019). Alternatively, it has been proposed to interpret the topography of classifier weights from the decoding analysis, and e.g. submit them to source estimation. In this case, the weights have to be back-projected into activation patterns before the application of source estimation (Haufe et al., 2014). This back-projection method should be applied whenever the classifier patterns themselves (rather than decoding accuracy) are interpreted, since otherwise seemingly high contributions from some sensors or voxel may be due to noise rather than signal (Haufe et al., 2014). This offers another approach to decoding analysis in source space: One can apply decoding to data across all ROIs that are assumed to be involved in the processes of interest, and interpret the distribution of these decoding weights across ROIs following back-projection, thus determining which regions contribute more or less to the successful decoding of stimulus or task information. To our knowledge, these two approaches – conventional ROI-by-ROI decoding and across-ROI decoding with back projection – have not been applied to the same EEG/MEG dataset in source space yet.

The anterior temporal lobes are sensitive to the semantic characteristics of single words as shown in several EEG/MEG studies both in univariate analyses (Dhond et al., 2007; Farahibozorg et al., 2022; Marinkovic et al., 2003) as well as in multivariate analyses on intracranial recordings (Chan et al., 2011; Rogers et al., 2021).

A recent study examined the activation dynamics in several brain areas within the semantic network by contrasting two different tasks that differed with respect to the depth of semantic processing: semantic decision (SD) and lexical decision (LD) (Rahimi et al., 2022). The evoked responses indicated that different semantic task demands are associated with an early modulation of visual and attentional processes, followed by differences in the semantic information retrieval in the ATLs, and finally a modulation in control regions (PTC and IFG) involved in the extraction of task-relevant features for response selection. These results were corroborated by the functional connectivity analysis in this and another study that revealed connectivity between the ATLs and semantic control regions (Rahimi et al., 2022; Rahimi, Jackson, Farahibozorg, et al., 2023).

The present study aimed to extend the evoked results of Rahimi et al. (2022) using a multivariate decoding approach in source space. In contrast to the previous study, we investigated both stimulus and task features. In addition, we evaluated two different decoding approaches: per-ROI and across-ROI. We used these methods to compare the spatio-temporal decodability of stimulus and task features of dynamic brain activation patterns. In particular, we asked 1) whether the patterns that carry stimulus and task information overlap in space and time, and to what extent this information is distributed vs. localised; and 2) whether task affects the amount of available stimulus-specific information within the semantic network.

## Methods

We used the dataset described in Farahibozorg (2018) and Rahimi et al. (2022), and most pre-processing steps are identical to that study. Thus, we will here report a summary of the most relevant characteristics and refer to the previous study for more detailed information. Previous analyses of this dataset did not employ decoding of task or stimulus features. The code for the analysis is available on https://github.com/magna-fede/SourceSpaceDecoding_SDvsLD.

### Participants

We analysed data from 18 participants. All of them were native English speakers, right-handed, with normal or corrected-to-normal vision, and reported no history of neurological disorders or dyslexia. The study was approved by the Cambridge Psychology Research Ethics Committee.

### Stimuli and Procedure

250 uninflected words were used in the visual stimulation. Each word belonged to one of 5 different categories based on its semantic content: visual, auditory, hand-actions, emotional or neutral abstract words. Table 1 summarizes the psycholinguistic variables. For the stimulus feature decoding, we used only a subset of the stimuli. Specifically, we created a subset of the two abstract words categories (i.e., emotional and neutral categories), so that we ended up with 4 semantic categories with 50 stimuli each. This was done because the stimuli in the emotional and neutral categories contained 1.5 letters more on average, compared to the other three categories. To avoid a false semantic classification due to word length, we removed this confound by selecting 50 words across these two categories so to have a better match for possible confounds, resulting in a more generic abstract word category.

**Table 1.** Summary statistics (mean and (standard deviation)) for each of the (single-word) semantic categories presented in both the SD and LD task, and for pseudowords presented only in the LD task.

|  | Number of<br>letters | Frequency | Orthographic<br>Neighbourhood<br>Size | Bigram<br>Frequency | Trigram<br>Frequency |
| --- | --- | --- | --- | --- | --- |
| **Action** | 5.02<br>(1.20) | 19.67<br>(30.86) | 5.44<br>(5.25) | 17368.95<br>(9580.53) | 1690.77<br>(2505.16) |
| **Auditory** | 5.16<br>(1.22) | 11.03<br>(17.25) | 5.28<br>(5.24) | 17599.13<br>(9181.36) | 1825.17<br>(2426.59) |
| **Visual** | 5.18<br>(1.48) | 13.32<br>(17.58) | 4.50<br>(5.44) | 20089.52<br>(11698.66) | 1867.21<br>(3314.68) |
| **Abstract** | 5.28<br>(0.93) | 23.64<br>(23.76) | 3.26<br>(3.84) | 19495.50<br>(9564.39) | 1597.65<br>(955.62) |
| **Pseudowords** | 5.00<br>(1.00) | N/A | 4.70<br>(4.50) | 19465.83<br>(10435.78) | 1670.66<br>(2029.80) |

The study consisted of four blocks and the full set of word stimuli was presented in each of them. One block consisted of a lexical decision (LD) task, where participants were presented with 250 words and an additional 250 pseudowords. They were asked to press a button with either their left index or ring finger to indicate whether the letter string represented a real, meaningful word or not. The other three blocks were all different instances of a semantic decision (SD) task: Participants were presented with the same 250 words as for LD, plus 50 additional fillers and 30 target words. Participants had to press a button with their left middle finger whenever the current stimulus belonged to the specific target semantic category for that block. The three target semantic categories were: (1) “food that contains milk, flour, or egg”, (2) “non-citrus fruits”, and (3) “something edible with a distinctive odour”. The target categories were unrelated to the word categories of the stimuli described above. The order of presentation of the SD blocks was randomised; half of the participants performed the LD task first, and half of them did it after the SD blocks.

In the SD block, only non-target trials were included in the analyses, i.e. trials that did not require a button press response. In the LD task participants were responding with a button press to both real words and pseudowords since a two-alternative forced choice is the standard procedure in LD tasks. We do not consider the details of response execution at the end of each trial as a serious confound for our EEG/MEG results in earlier latency ranges (see Rahimi et al., 2022 for details). These potential confounds are present only when classifying task in SD vs LD blocks, but not when considering the classification of different SD blocks or when performing single-word semantic classification.

### Data acquisition and source estimation

Simultaneous EEG and MEG data were concurrently recorded to maximize spatial resolution (Molins et al., 2008). The sampling rate during data acquisition was 1000 Hz and an online bandpass filter 0.03 to 330 Hz was applied. Signal Space Separation with its spatiotemporal extension as implemented in the Neuromag Maxwell-Filter software was applied to the raw MEG data. Raw data were visually inspected for each participant, and bad EEG channels were marked and linearly interpolated. Data were then band-pass filtered between 0.1 and 45 Hz. FastICA algorithm was applied to the filtered data to remove eye movement and heartbeat artefacts. We used L2-Minimum Norm Estimation (MNE) for source reconstruction (Hämäläinen & Ilmoniemi, 1994; Hauk, 2004). Three-layer boundary element forward models were constructed using structural MRI scans.

### Regions of interest

We focused our analyses on six regions of interest that were defined using the anatomical masks provided by the Human Connectome Project (HCP) parcellation (Glasser et al., 2016). We adhered to Rahimi’s et al. (2022) selection, which examined the evoked responses and functional connectivity analysis of these same ROIs (Figure 1). These areas were chosen due to their putative role in the semantic brain network, as derived from neuroimaging fMRI studies. ATLs have been proposed to be multimodal semantic hub regions (Lambon Ralph et al., 2017). IFG (Inferior Frontal Gyrus) and PTC (Posterior Temporal Cortex) have been described as semantic control regions required for the appropriate task- or context-relevant processing of a word stimulus (Jackson, 2021). Angular Gyrus (AG) has been suggested as an additional hub region or convergence zone (Binder, 2016). Finally, we included PVA (Primary Visual Areas) to test potential task effects on early perceptual processing.

**Figure 1.**
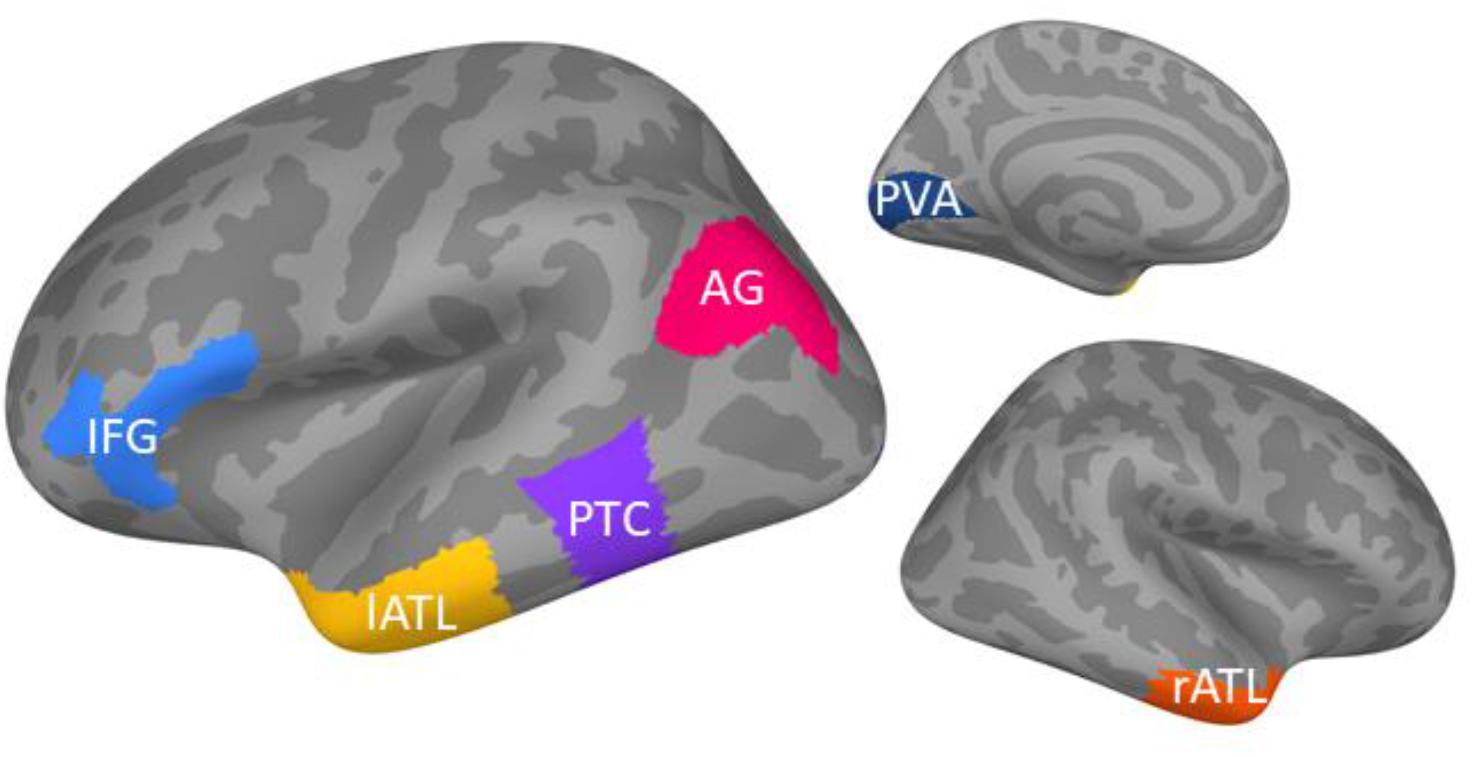
Regions of interest (ROIs) used in this study (based on Rahimi et al. (2022)).

### Pre-processing

The pre-processing and decoding analyses are based on tutorials and examples available on MNE-Python (Gramfort et al., 2013) and on the tutorial provided by Grootswagers et al. (2017). We estimated brain responses in source space for each participant and for each trial. We sampled our data in epochs that started from 300 ms before the stimulus onset until 900 ms post-stimulus. Data was down-sampled to 250 Hz. For the decoding analysis, we intended to keep as much information in our activation patterns as possible and therefore used signed source estimates, i.e. rather than taking the (only positive-valued) intensities of dipole sources at each vertex, we kept their directions of current flow. We present the grand-averaged time courses of brain activation for each ROI and for different tasks in Figure 2 (using the “mean-flip” option to account for the variability of source orientations). The figures show that we obtained clear evoked responses in all ROIs and conditions.

**Figure 2.**
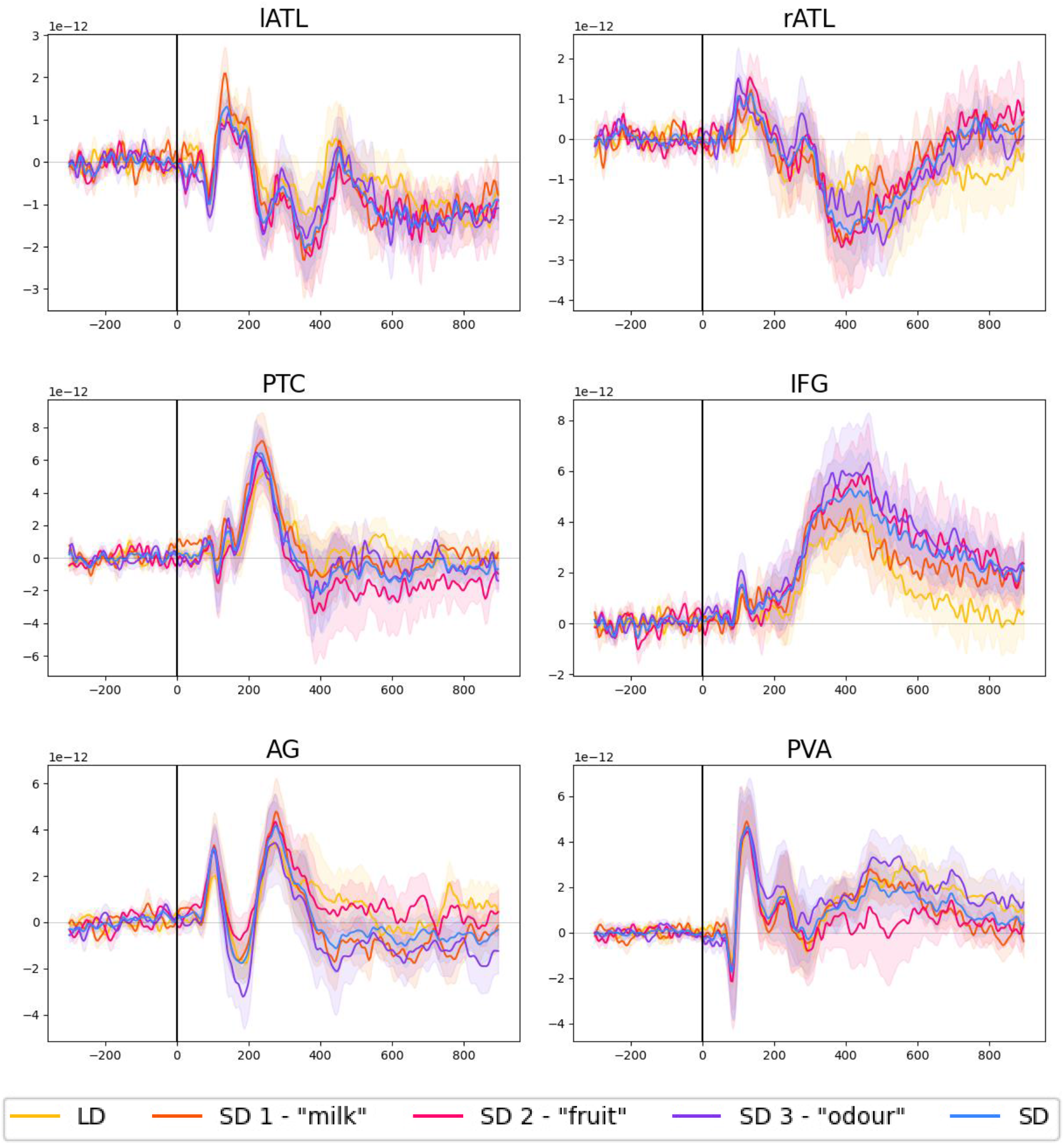
Evoked responses for each task and semantic decision tasks averaged together. Shadowed area report standard error of the mean (LD: lexical decision; SD: semantic decision; “milk”, “fruit” and “odour” decision refer to single SD blocks).

The pre-processing consisted in averaging 3 trials together for each class, in order to reduce noise in the data and improve decoding accuracy (Isik et al., 2014). We then z-score-normalised each of the resulting epochs. The models fitted were always logistic regression models, using a five-fold cross-validation procedure. We calculated accuracy as the average Receiver Operator Characteristic (ROC) Area-Under-Curve (AUC) on the test data across the 5 folds.

### Decoding Analysis

We first decoded task in two different ways: LD-SD, where we classified whether a trial belonged to a lexical decision or semantic decision block; and SD-SD, where we decoded to which of the 3 SD blocks a certain trial belonged. Then, we decoded single-word semantic category, separately for LD and SD blocks. For each classification analysis, we compared two different decoding approaches: A) Individual ROIs: we estimated decoding accuracy in each ROI separately; B) Combined ROIs: we fitted a model that used all vertices across all ROIs, then determined the contribution of each ROI by looking at the root-mean-squared of the back-projected weights of the model within ROIs.

#### Individual ROIs Accuracy

In the LD-SD classification, we separately classified each of the LD-SD combinations using a binary logistic regression model, and accuracy was calculated as the average across them. In the SD-SD and stimulus feature classification we fitted multinomial logistic regression models and calculated accuracies as one-vs-rest ROC AUC (this ensured that the chance level was 0.5 for all tasks). For the semantic category analysis, we fitted a multiclass logistic regression model. For SD blocks, we estimated accuracy both when concatenating the 3 blocks in one model (in order to have more trials, and therefore a more reliable estimate of regions and time points where stimulus features were decodable) as well as separately per block (to have a fair comparison in terms of noise level with the LD block with a comparable number of trials).

We performed statistical significance analyses to test differences in accuracy between different classification tasks, specifically, we tested A) whether task decoding accuracy differed between LD-SD and SD-SD; B) whether we could reliably decode word category classification above chance; C) whether stimulus feature decoding accuracy differed between LD and SD tasks. We used a cluster-based permutation test over time to correct for multiple comparisons (Maris & Oostenveld, 2007).

#### Combined ROIs Patterns

In this analysis, we decoded patterns combined across ROIs and then determined how much each ROI contributes to the corresponding classifier. For this purpose, we back-projected the classifier weights into the corresponding activation patterns across ROIs as suggested by Haufe et al. (2014). This back-projection is required because the classifier weights may reflect both signal and noise, while the back-projected patterns reflect brain activity predicted by the estimated model.

In general, we followed the same steps as in the individual ROIs procedure described in the previous section, and subsequently extracted the back-projected weights of each feature in the model, which we will refer to as activation patterns or just patterns from now on.

For each classification task, we will report the root-mean-squared (RMS) activation patterns for each ROI, averaged across participants. As each model had a set of weights specific to each class, we will first calculate an ROI’s RMS of each class-specific activation pattern – for example in the semantic classification, ROI-specific hand-class patterns (and analogously for visual, auditory, and abstract classes). Then, by averaging across classes, we obtain the average activation pattern time course of each ROI, for each participant. The RMS pattern is a measure of the relative contribution of a certain ROI to the classification: Each vertex’s value reflects the relative contribution of that specific vertex to the signal used by the classifier, i.e. vertices that are close to 0 are relatively less informative, whereas larger absolute values indicate that the vertex contains more information. By considering the RMS instead of the average, we avoid that patterns of vertices within the same ROI with different polarities will cancel each other.

## Results

In the following, we will present results for different task classifications (LD-SD and SD-SD; Figure 3) and semantic word category classification (Figure 4). For each part, we will first present the results for individual-ROI classification followed by combined-ROI classification.

**Figure 3.**
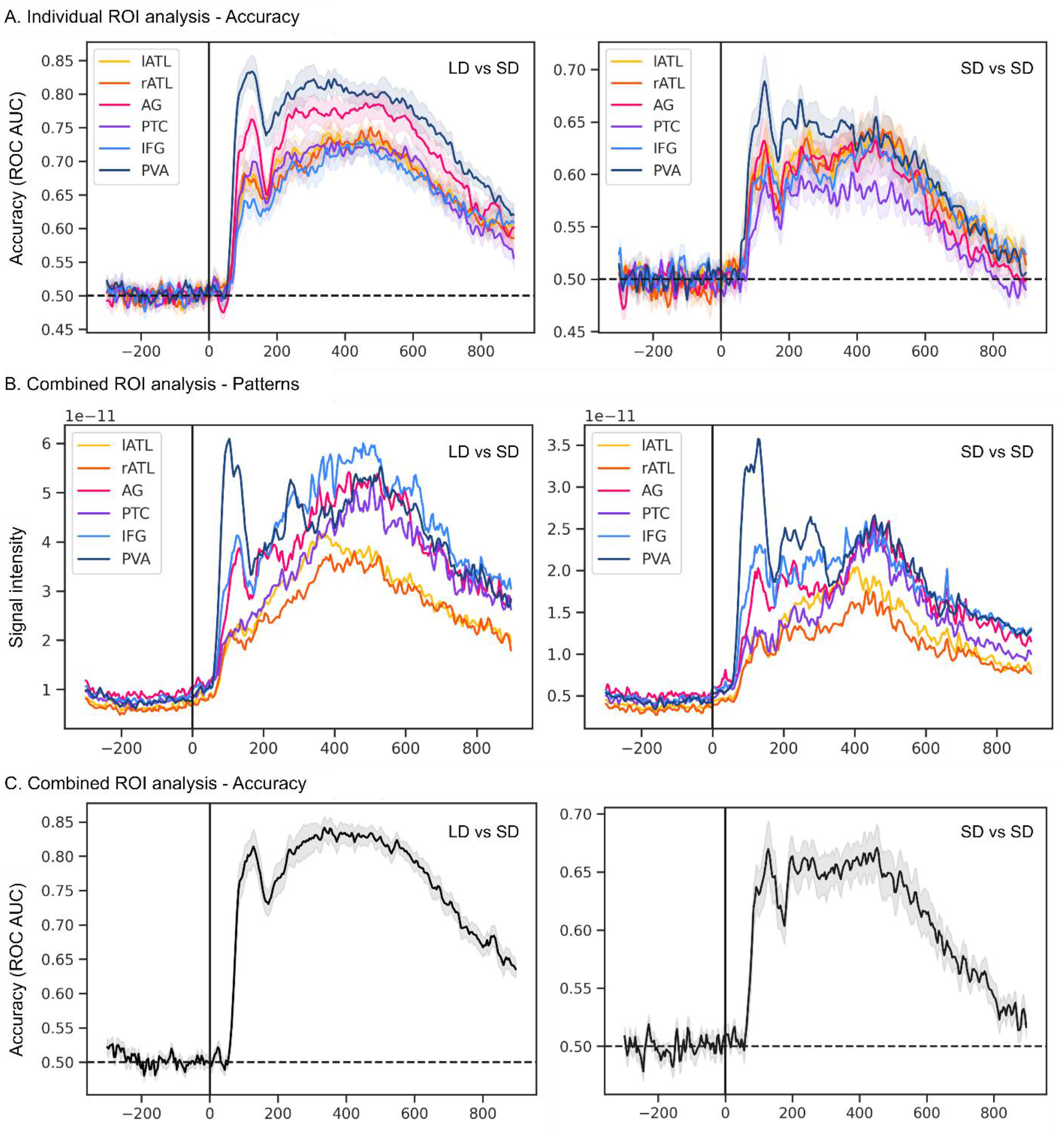
Task classification performance. For all plots, the shadowed area represents the standard error of the mean (across participants). **A**. Individual ROI task classification accuracy (ROC AUC) when decoding LD from SD (left) and when decoding SD tasks (right). **B**. Root-mean-square of the activation patterns within each ROI, when decoding LD from SD (left) and when decoding SD tasks (right). **C**. Combined ROI task classification accuracy (ROC AUC) when decoding LD from SD (left) and when decoding SD tasks (right).

**Figure 4.**
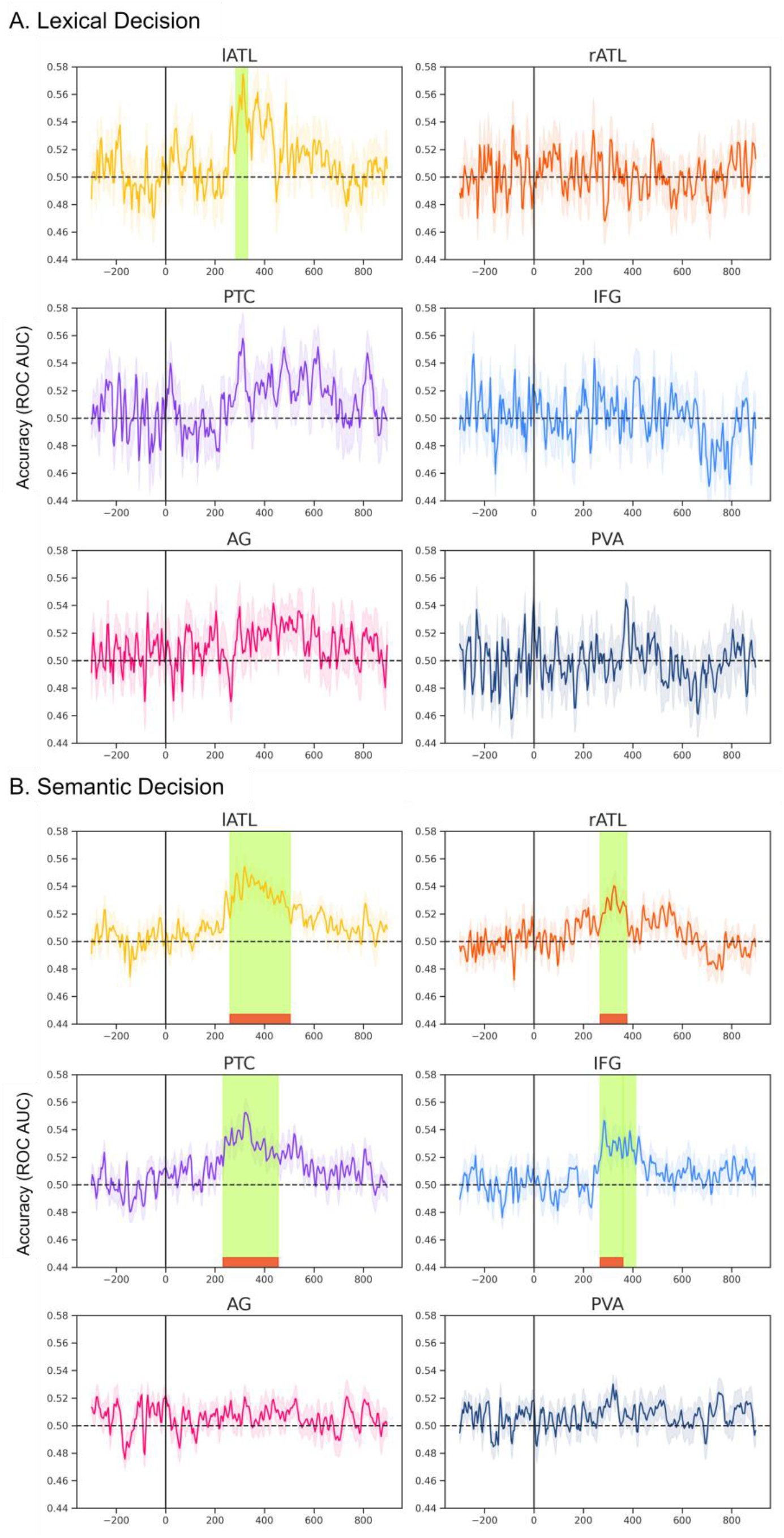
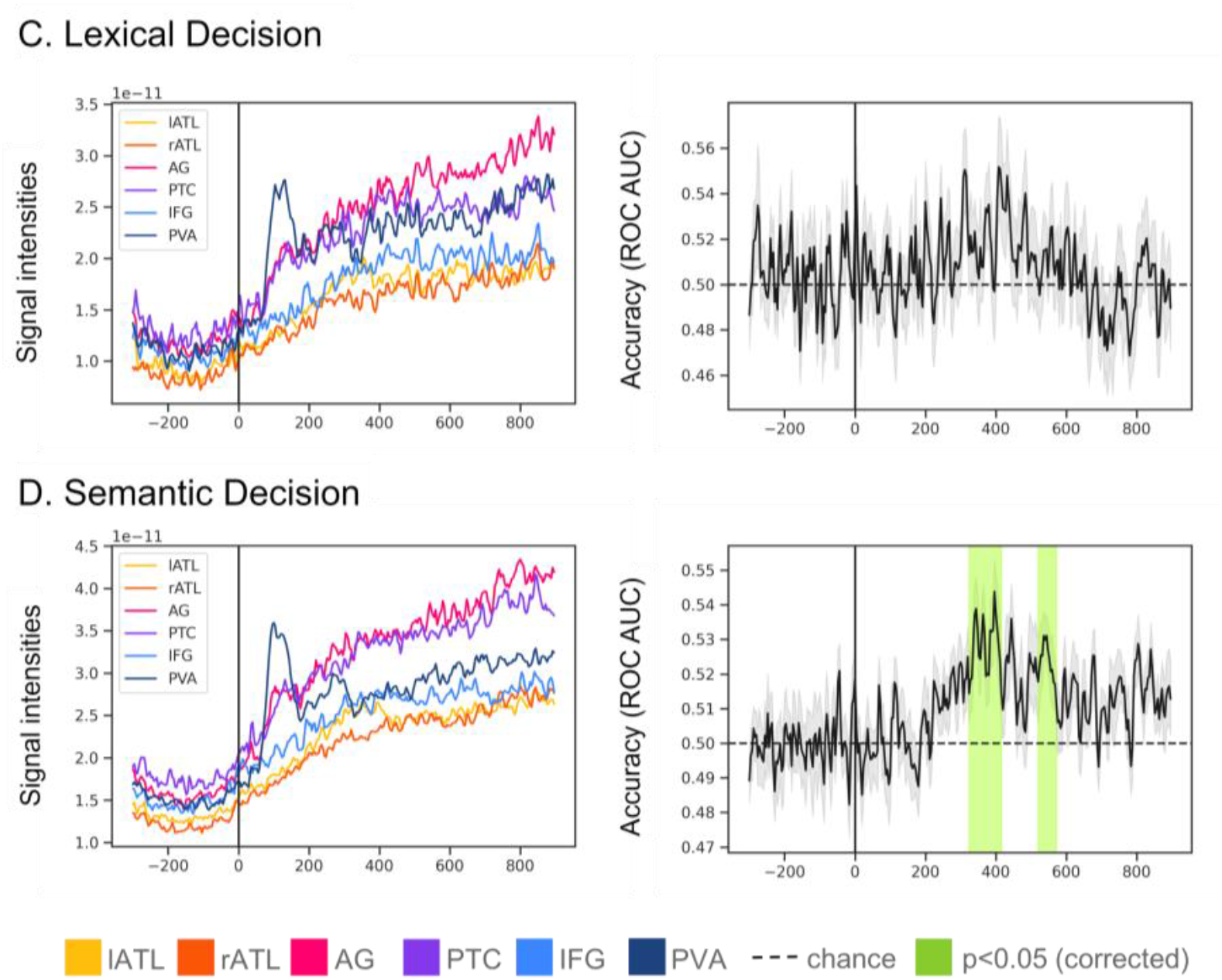
Single-word semantic category classification performance. For all plots, the shadowed area represents the standard error of the mean (across participants). Green colour highlights the times when cluster-based permutation correction test revealed a significant cluster (α=0.05) where accuracy was above 0.5, and red the clusters that are still significant after Bonferroni correction (across ROIs). **A**. Individual ROI task classification accuracy (ROC AUC) when decoding semantic features in LD block. **B**. Individual ROI task classification accuracy (ROC AUC) when decoding semantic features in SD blocks. **C**. Combined ROI root-mean-square of the activation patterns within each ROI (left) and classification accuracy (ROC AUC) (right), when decoding semantic features in LD block when decoding LD. **D**. Combined ROI root-mean-square of the activation patterns within each ROI (left) and classification accuracy (ROC AUC) (right), when decoding semantic features in SD block.

### Task classification

Figure 3 shows classification accuracy results for the task comparisons (left: LD-SD, right: SD-SD) over time for different ROIs in source space, separately for the individual-ROI approach (A) and the combined-ROI approach (B and C).

Both task classifications show similar results, in so far as task was decodable from before 100ms after stimulus presentation and then for most of the epoch, both in the individual- and combined-ROI approach. Detailed information is presented below.

#### Individual ROIs Accuracy

In LD-SD, accuracy is at chance level in the baseline period (before stimulus presentation), but it sharply increases after approximately 50 ms post-stimulus reaching accuracies of over 0.7 in all examined ROIs at around 100 ms. Accuracy plateaus for a large part of the epoch and is followed by a slow decline. The SD-SD classification shows very similar results, although accuracy is generally lower, peaking below 0.7. Cluster-based permutation testing (not shown) confirms this difference with LD-SD accuracy being significantly higher than SD-SD in all ROIs from around 70 ms post-stimulus presentation to the end of the epoch (apart from IFG, where the difference starts at approximately 152 ms).

#### Combined ROIs Patterns

When combining all ROIs in one model, the accuracy profile is very similar to that in the individual-ROI approach, i.e. it is characterized by a sharp increase soon after stimulus onset and a plateau afterwards (Figure 3C). However, while in the previous analysis the shape of accuracy time courses was very similar across ROIs, we observed larger differences among ROIs when examining the RMS of activation patterns within ROIs (Figure 3B). For example, PVA’s (dark blue line) contribution to task classification (both LD-SD and SD-SD) peaks around 100ms. AG’s and IFG’s time courses (pink and light blue, respectively) are similar, with a smaller peak than PVA around 100ms, but similar and even larger values after 250 ms. In contrast, the temporal lobes (lATL and rATL, yellow and orange) and posterior temporal cortex (PTC, purple) show similar time courses without a distinct early peak. Their activations slowly increase until they reach their peak around 400 ms. Also in this case, cluster-based permutation testing (not shown) showed that decoding accuracy was higher in LD-SD and SD-SD from approximately 64 ms post-stimulus onset.

Thus, while the Individual-ROI analyses revealed that all ROIs carry information about the tasks, the Combined-ROI provides more information about the time courses of information, in particular about their differential contributions at early (∼100ms) versus later (>250ms) latencies.

### Single-word semantic features classification

In the semantic category classification, for each stimulus we decoded the semantic category it belonged to (out of 4 alternatives), separately for LD and SD blocks. Figures 4A and 4B show results for the individual ROIs stimulus feature decoding. Figures 4C and 4D show results for the combined ROIs pattern and accuracy. In general, we observed that stimulus features were decodable above chance level when participants were engaged in a semantic decision task, but less so in lexical decision.

#### Individual ROIs Accuracy

Figure 4A revealed that in LD blocks only lATL showed significant decoding for word category and only in one short time window around 300 ms. In contrast, for SD we obtained significant decoding in several regions (left and right ATL, PTC and IFG). SD decoding survived Bonferroni correction across ROIs. For each ROI we report the significant cluster(s) and their approximate latencies in Table 2. In SD, we observed above-chance decoding accuracy in left and right ATL, PTC and IFG, but not in PVA and AG. PTC successfully classifies single-word semantic category between 236-452 ms post-stimulus. This is followed by left and right ATL, respectively, between 264-500 and 272-372 ms. IFG was above chance approximately at the same time, between 282-356 and 364-408 ms (but the second cluster did not survive Bonferroni correction). As the SD model was trained on trials from 3 blocks, the table reports also the results of cluster-based permutation tests for each SD block separately, to show results from a comparable noise level to the single LD block. Decoding accuracy in single SD blocks was above chance in more ROIs-latency combinations compared to the LD block (apart from the odour task).

**Table 2.**
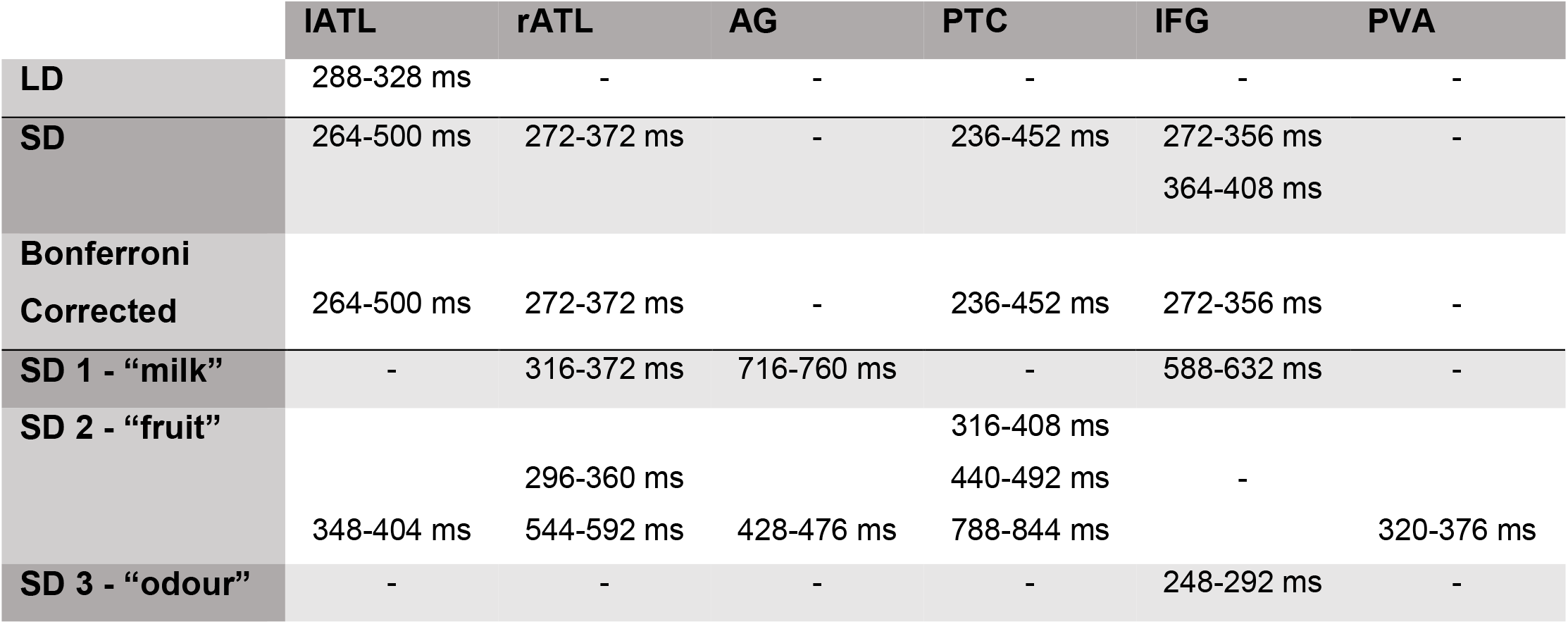
Approximate timings when single-word semantic category is decodable in each ROI. SD refers to the information when fitting a model to all SD blocks concatenated, whereas the effects for models fitted to individual SD blocks (i.e., (1) “milk”, (2) “fruit”, and (3) “odour”) are reported in the last three rows of the table.

We then statistically tested whether the decoding accuracy was significantly higher in SD than in LD considering that word category was decodable only in lATL in the LD block, in contrast to all the semantic network in SD. However, no ROI showed a significant effect. To further understand whether task affected the stimulus category decoding, we computed cross-task decodability. This analysis was included to determine whether the representation of specific semantic information was consistent across tasks (i.e., successful cross-task decoding indicates that multivariate patterns of a certain word category are represented similarly irrespective of tasks). We trained a model on SD trials and tested stimulus features decodability of the same model in LD trials, and vice versa. Results are reported in Figure 5. When the model was trained on SD and tested on LD trials, we observed above-chance accuracy in all semantic areas: lATL between 276-408 ms, rATL between 240-456 ms, PTC between 272-512 ms, IFG between 360-408 ms, and PVA between 184-240 ms (where the last two regions did not survive Bonferroni correction across ROIs). We observed similar results in models trained on LD and tested on SD (although LD trials were significantly less so we will not interpret the results for each individual ROI).

**Figure 5.**
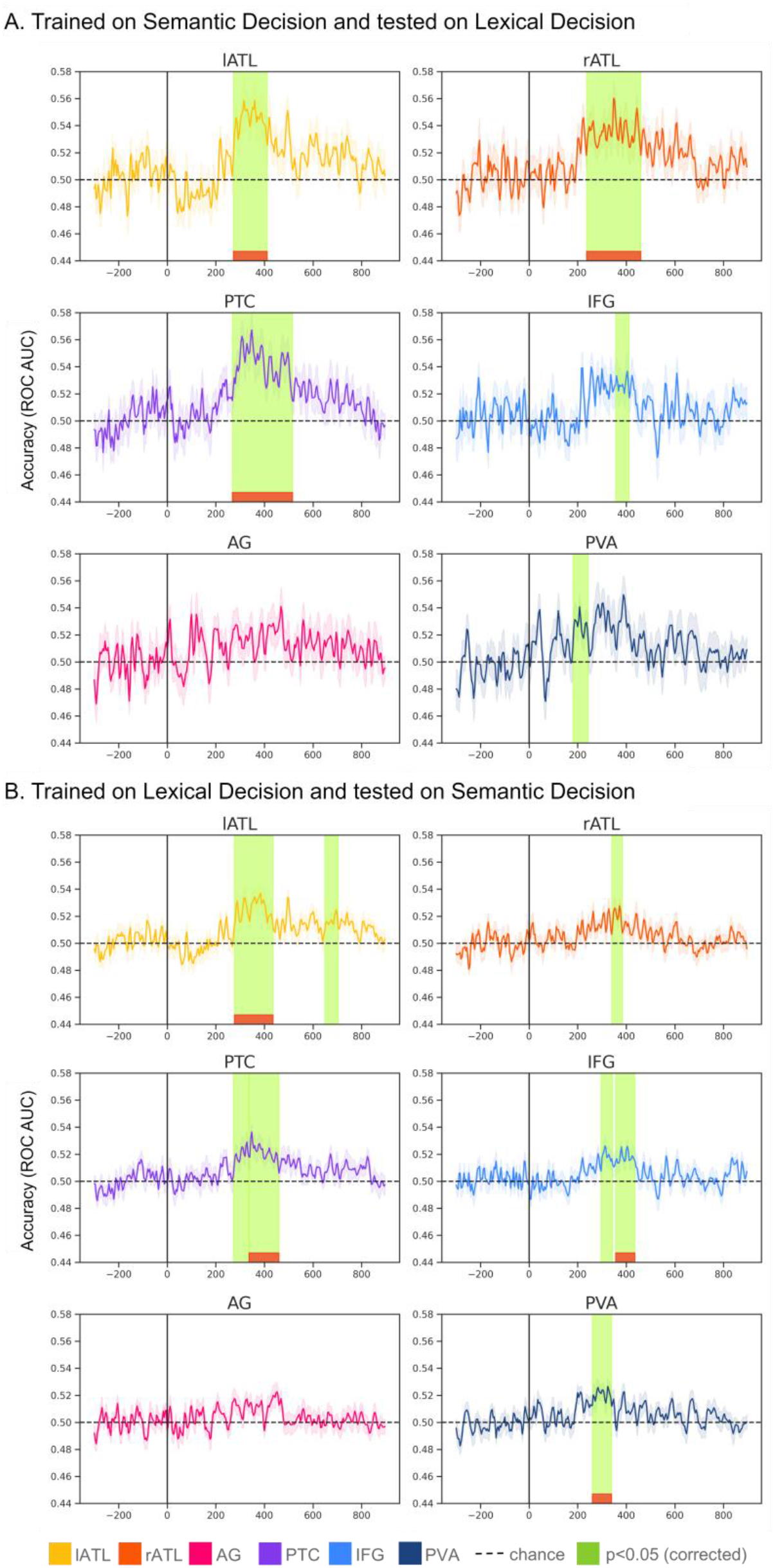
Cross-task semantic category decodability. For all plots, the shadowed area represents the standard error of the mean (across participants). Time points highlighted in green are times when cluster-based permutation correction test revealed a significant cluster (α=0.05). In red, are clusters that are still significant after Bonferroni correction (across ROIs). **A**. Individual ROIs cross-decoding when the model was trained on SD trials and tested on LD trials **B**. Individual ROIs cross-decoding when the model was trained on LD trials and tested on SD trials.

As a final step, we wanted to determine whether the decoding performance was driven by any particular semantic word categories, so we examined the confusion matrices of each significant temporal cluster. To obtain a confusion matrix for each cluster, we obtained the model’s predicted class for each time point, separately for each participant. We then fitted a (normalised) confusion matrix for each point and considered the average values across time for each cluster. Figure 6 shows the confusion matrices for all the temporal clusters that were significantly above chance across participants in the semantic category classification, for each task/ROI separately. The confusion matrices indicate that abstract words are the most differentiable (i.e., relatively higher accuracy) from the other classes, which were all concrete.

**Figure 6.**
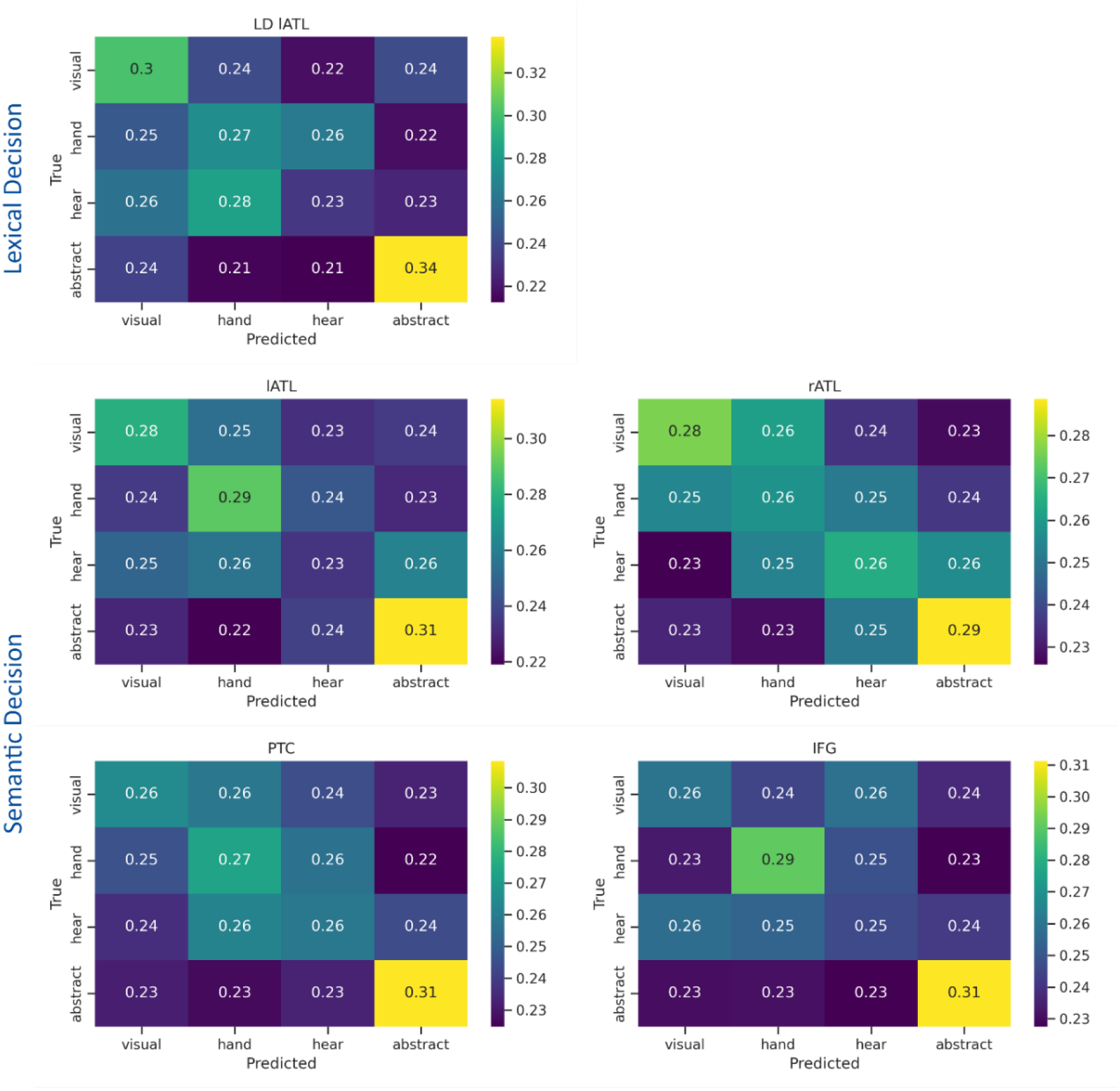
Confusion Matrices for the decoding of semantic word categories. Each cell represents the temporally-averaged probability of a predicted word category, given the true word category, separately for each of the significant temporal clusters found in the semantic category classification. Smaller values indicate lower probabilities. In all cells, abstract words were the most accurately predicted category, compared to the other (concrete) words.

#### Combined ROIs Patterns

Similarly to the individual ROI case, accuracy was reliably above chance only in the SD but not LD task. In SD, one cluster extended between 328-412ms and a second cluster approximately between 524-568ms. However, also in this case, statistical analysis revealed no difference between SD and LD models. Decoding accuracy for individual ROIs as well as for combined ROIs was quite low (below 0.55) compared to our results in Figure 3. As a result, the activation patterns for the Combined-ROI analysis appeared to be less informative in this case. For both LD and SD these patterns showed an almost linear, drift-like increase over time following the baseline period, with a small peak at about 100ms for PVA. This confirms the assertion of Haufe et al. (2014) that classifier weights and activation patterns are less interpretable when decoding accuracies are low.

## Discussion

Our study had two main goals: First, we wanted to characterise the spatiotemporal dynamics within the semantic brain network using multivariate information decoding; and second, we compared two different approaches to source space decoding, based on individual ROI and combined ROI data.

Logistic regression models performed very well at classifying task and showed high accuracy in decoding LD vs SD trials in all ROIs from before 100ms. This is consistent with previous studies that reported effects of task on visual word processing from early latencies (around 100ms, Chen et al., 2015) and the evoked and functional connectivity results obtained from the same data (Rahimi et al., 2022). However, these effects between LD and SD may not be surprising, since these tasks differ in several ways, such as the semantic depth of processing, task difficulty and attentional demands (as reflected in response times), and stimuli were presented in different blocks. More interestingly, the classifier could distinguish with high accuracy between different SD blocks, which reflected more subtle task demands as they only differed with respect to the semantic target category for the response. In all task classifications, all ROIs showed an early increase in decoding accuracy from about 50 ms which was followed by a plateau, and a slow decline towards the end of the epoch. This plateau indicates that for a sustained period, the whole semantic network carries information about the task that is being performed.

Also, we were able to decode single-word semantic category across the whole semantic network (left and right Anterior Temporal Lobe, left Posterior Temporal Cortex, and left Inferior Frontal Gyrus) in the semantic decision task, and in the left Anterior Temporal Lobe in the lexical decision task. This indicates that semantic information is represented across all the semantic network, at least when performing a semantically engaging task. However, statistical tests revealed no significant differences between the two tasks in the direct comparison in any ROI. Thus, word category information may be present across the semantic network in both tasks, but only reach significance in the SD task. This was confirmed by our additional cross-task decoding analysis: when training the classifier on semantic decision and testing on lexical decision trials, and vice versa, we could successfully decode stimulus features to a similar degree as in individual tasks. Thus, while a previous study revealed significant differences between tasks in evoked activity and functional connectivity (Rahimi et al., 2022), our current decoding results show that when it comes to semantic feature representation there is a higher degree of similarity across tasks. Altogether, this supports the view that early visual word recognition processes are flexible, i.e. they are modulated by task demands, but semantic information is processed even if not explicitly required by the task (Chen et al., 2015; Evans et al., 2012).

Only Angular Gyrus and Primary Visual Cortex did not produce above-chance decoding accuracy for semantic word category, neither in LD nor in SD blocks. This confirms previous results that early brain activity in the temporal lobes, but not in angular gyrus, reflects semantic word features (Farahibozorg et al., 2022). It also supports the view that AG does not support semantic representation per se, but is involved in more spatiotemporally extended semantic processing (Humphreys et al., 2021). The absence of PVA effects in the category decoding analysis indicates that perceptual properties of the stimuli were not influencing the classification performance, as our stimuli were well matched for several relevant psycholinguistic variables. Thus, while for pictorial stimuli semantic category is often confounded by visual stimulus features (e.g., natural concepts being more “curvy” and artefacts being more “edgy”), our word recognition results are most likely due to semantic stimulus features.

Finally, we tested whether and how different semantic features affected the decoding performance in different regions. We addressed this question by computing confusion matrices for the decoding of different word categories against each other. This revealed that concrete word categories were more likely to be confused with each other (mostly indistinguishable) and that abstract words were driving the decoding performance across all regions that showed above-chance performance (i.e., all the semantic network in the SD task). This confirms that information about a word’s concreteness is represented in the activation patterns within the semantic brain network from early latencies, as demonstrated in previous studies at a univariate level (e.g., Farahibozorg et al., 2022). Also, this indicates that multivariate brain patterns reflect semantic similarities (i.e., more similar concepts have similar representations) and parallel fMRI results (Kriegeskorte et al., 2008).

A methodological objective of our study was to compare two approaches to source-space decoding. We looked at both task classification and stimulus classification using decoding per ROI and decoding across ROI. While we found reliable task decoding using both approaches, for the semantic category decoding only the individual-ROI approach yielded reliable results. This suggests that when the effect of interest is distributed across brain regions, then the across-ROI approach is more informative, but when the effect of interest is distributed more sparsely over space and time, i.e. if it is present only at certain latencies or regions, then the per-ROI approach is more sensitive.

Decoding performance was high both for general (LD-SD) and more subtle (SD-SD) tasks, across all our ROIs. Although in both cases accuracy varied across ROIs (e.g., relatively higher in PVA and AG compared to semantic ROIs in the LD-SD task), it is not straightforward how to interpret this as evidence of different degrees of information contained in each ROI, as different algorithms and parameters will likely influence the accuracy score. For example, this difference could just reflect the “visibility” of a region by the EEG/MEG sensors. Hence, the individual-ROI approach was not very informative concerning the time course of the effects (contrary to the semantic category decoding), as all ROIs showed a similar timecourse (rapid increase in performance, followed by a plateau, and slow decline). However, more information was available when observing the activation patterns of the model that combined across-ROIs information. For example, we observed that while the relative contribution of visual areas starts early, semantic regions take longer to reach their peak, consistent with univariate results of a posterior-to-anterior sweep of information which is observed in evoked responses (Marinkovic et al., 2003; Rahimi et al., 2022). Interestingly, we found no obvious difference between ROIs relevant for the LD-SD and SD-SD classifications.

The only region that showed significant decoding accuracy across all our analyses was the left ATL, with the caveat that the differences in stimulus decoding accuracy did not differ statistically between the lexical and semantic decision task. Nevertheless, we found involvement of those regions that have been put forward as the core semantic network (Lambon Ralph et al., 2017). More specifically, it has been suggested that IFG exerts semantic control function via PTC onto ATL (Jackson et al., 2021). Our results are consistent with this framework. Information regarding the semantic properties of a word seems to be spread across the core semantic network, but probably not outside (at least not in PVA and AG). This semantic information is at least partially stable across tasks. Future studies should investigate the information flow based on multivariate pattern as well as connectivity analyses in more detail. The amount of multivariate methods available for evaluating semantic computations from neuroimaging data is rapidly increasing (see Frisby et al., 2023 for a review). Furthermore, methods that characterise brain connectivity based on multidimensional relationships, or pattern-to-pattern transformations, have recently become available (Basti et al., 2020; Rahimi, Jackson, Farahibozorg, et al., 2023; Rahimi, Jackson, & Hauk, 2023).

In conclusion, our results demonstrate that EEG/MEG source space activity contains rich information about stimulus and task features in written word recognition, which will be essential to unravel the complex dynamics in the semantic brain network.

## Acknowledgements

This work was supported by intramural funding from the Medical Research Council UK (MC_UU_00005/18), and by Cambridge ESRC DTP and Cambridge European & Newnham College scholarships awarded to F.M. For the purpose of open access, the author has applied a CC BY public copy-right license to any Author Accepted Manuscript version arising from this submission. We thank Rezvan Farahibozorg for data collection and information on the dataset. We would like to thank also Setareh Rahimi, Pranay Yadav, and Kevin Campion for helpful discussions and support during the preparation of this manuscript.

## Notes

### Competing Interest Statement

The authors have declared no competing interest.

### Summary of Updates

Updated acknowledgements and references; improved figure visualisation; included code availability statement; copy-editing.

